# Biophysical Characterization of an Essential Mammalian Protein; Transcription Termination Factor I (TTF1)

**DOI:** 10.1101/2022.08.20.504633

**Authors:** Kumud Tiwari, Gajender Singh, Samarendra Kumar Singh

## Abstract

Mammalian Transcription Terminator Factor 1 (TTF1) is an essential protein which plays diverse cellular physiological functions like transcription regulation (both initiation and termination), replication fork blockage, chromatin remodelling, DNA damage repair etc. Hence, understanding the structure and mechanism conferred by its variable confirmations becomes significantly important. But so far, almost nothing is known about the structure of either the full-length protein or any of its domain in isolation. Moving towards achieving the above goals, our lab has codon optimised, expressed and purified N-terminal 190 amino acid deleted TTF1 (ΔN190TTF1) protein, since full length protein even after multiple trials could not be purified in soluble form. In this article, we have characterized this essential protein by studying its homogeneity, molecular size and secondary structure using tools like dynamic light scattering (DLS), circular dichroism (CD) spectroscopy, Raman spectroscopy and atomic force microscopy (AFM). By CD and DLS we have shown that the purified protein is homogenous and soluble. CD spectroscopy also revealed that ΔN190TTF1 is a helical protein which was further confirmed by analysis of Raman spectra and Amide I region deconvolution studies. AFM imaging data discovered the size of single protein molecule to be 94 nm which is in agreement with the size determined by the DLS study as well. Our structural and biophysical characterization of this essential protein will open avenues towards solving the structure to atomic resolution and also will encourage the research to investigate the mechanism behind its diverse functions attributed to its various domains.

## Introduction

Eukaryotic DNA replication and transcription are vital processes occurring in three prime stages: initiation, elongation, and termination. Regulation of these stages are critically important specially during S phase of the cell cycle when certain regions of chromosomal DNA (like ribosomal DNA) are often transcribed and replicated simultaneously. Therefore, process of termination (polar fork arrest) is physiologically necessary to avoid collision between replication and transcription machinery which could lead to genomic instability and cell transformation. Polar fork arrest in mammalian cells takes place at certain defined regions of the chromosomes and among such regions are non-transcribed spacer region of ribosomal DNA (rDNA) flanking the area which codes for ribosomal protein. These particular sequences, situated 170 base pairs upstream and downstream of the rDNA gene repeats in mammalian cells are known as the Sal box elements (Figure 1a). In mammalian cells, these regulatory elements are responsible for site-specific Pol1transcription termination and DNA replication fork arrest in a polar fashion, which is mediated by the multifunctional protein; Transcription Termination Factor 1 (TTF1) upon binding with Sal box sequences^1^. Orthologues of TTF1 proteins have been found in a variety of organisms, including Rib2 of Xenopus, RNA Polymerase I enhancer binding protein (Reb1) of *S. cerevisiae* and *S. pombe*, and mitochondrial Transcription Termination Factor (mTERF) in human mitochondria^2,3^. Being a multifunctional protein, TTF1 is engaged in various cellular functions like chromatin remodelling, cell cycle regulation, DNA damage sensing, etc^4^. To execute these processes it also interacts with numerous regulatory proteins, including TTF1 Interacting Protein (TIP 5), Pol I and transcript release factor (PTRF), Mouse Double Minute 2 (MDM2), Cockayne Syndrome B (CSB), Alternative Reading Frame (ARF), etc. and catalyzes critical cellular functions in human^4–9^. These diverse activities make TTF1 truly a multifunctional and essential protein for the survival of the cell, misregulation of which has been linked with various cancers^1^–^12^.

**Figure 1.**
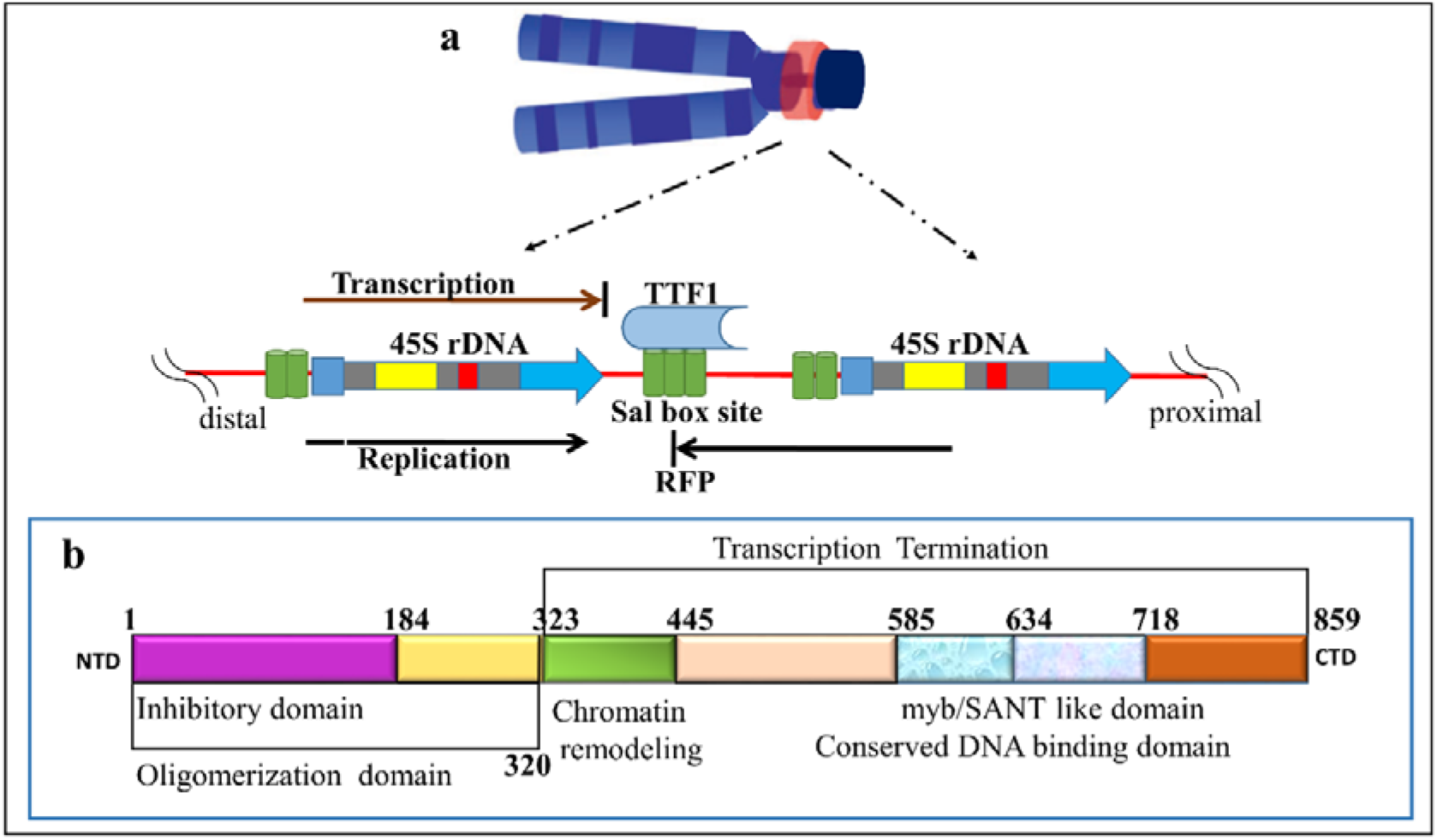
**a)**The diagram showing mouse chromosome de-condensed rDNA region [Nucleolar Organizing Region (NOR), red]; Schematic representation of the non-transcribed spacer region of rDNA showing rDNA genic region (45 S)^14^, Sal box site and fork pausing site RFP. TTF1 protein binds to Sal box and arrest replication from one direction (black arrow) and RNA pol1 mediated transcription from opposite direction (brown arrow)^1^; **b)**Domain architecture of mouseTTF1 protein showing N terminal inhibitory domain(1-184), oligomerization domain (1-320), chromatin remodeling domain (323-445), conserved DNA binding domain (myb/SANT like domain 585-718) and transcription termination domain (323-859)^13,15^.

Domain architecture of mouse TTF1 comprises a putative N-terminal regulatory domain, DNA binding trans-activation domain (myb/SANT domain) and C-terminal transcription termination domain. The N-terminal region of the protein, working as flap in oligomeric state, represses its DNA binding activity (Figure 1b)^13^.

However, to date, no research on its structural characterisation; neither computational nor physical has been reported yet. The very first article which reports ab-initio modelling of this essential protein has been communicated from our lab^16^. Considering the multifunctional and essential nature of TTF1, it becomes important to elucidate the structural knowledge in order to understand the mechanisms of action and use the same towards its therapeutic applications. Because multiple attempts to purify soluble full length TTF1 has been unsuccessful, we cloned codon optimised N terminal 190 amino acid truncated (ΔN190TTF1, to get rid of the inhibitory flap) mouse gene and expressed it in bacterial cells. In this article, we present biophysical and structural characterization of this essential protein using Dynamic Light Scattering (DLS), Circular Dichroism (CD) spectroscopy, Raman spectroscopy, and Atomic Force Microscopy (AFM) which will open up the field for further structural characterization of the protein to atomic level and make it available for understanding its mechanistic function in detail. This work would be the first ever report of biophysical and structural characterization of the essential mammalian protein TTF1.

## Materials and Methods

1. Sequence-based analyses: The amino acid sequence of mouse (*Mus musculus*) TTF1 protein was retrieved from UniProtKB (UniProtKB accession number: Q62187) and the physicochemical properties of TTF1 were analysed using ProtParam computational tool^17^. Gravy index score of proteins was measured by Kyte-Doolittle formula^18^.
2. Expression and purification of recombinant TTF1: Codon optimized N-terminal 190 amino acid (a.a.) truncated mouse TTF1 (ΔN190TTF1) gene was cloned in pMAL-c2X vector with N-terminal His tag and expressed in bacterial BL21-λDE3 strain. The tagged protein was then purified using Ni-NTA column. Further details of which are presented in the extended experimental procedures.
3. Dynamic Light Scattering (DLS): DLS analysis was performed in a Zetasizer Nano ZS (Malvern, Worcestershire, UK) equipped with 4mW He–Ne 633nm laser with a detection angle of 173° backscatter at 25°C, in a low volume (45uL) quartz cuvette. Measurements of intensity and volume in multiple narrow mode were used to derive the size distribution. Measurements were performed at least in triplicate with 15ug/ul protein in 25mM Tris-Cl pH 7.5 buffer. The dispersity (polydispersion index, Pdi) and the particles size were determined at 25°C performing four runs (5 measurements each).
4. CD spectroscopy: CD spectra was recorded using a Jasco J-1500 spectropolarimeter equipped with a Peltier thermos stated cell holder (Jasco, Easton, MD, US). Purified ΔN190TTF1 protein (2ug/ml) was diluted in Buffer A (15ug/ul in 25mM Tris-Cl pH 7.5 and 100 mM KCl). Data collection and analysis were carried out as described by Colarusso et al., (2018)^19^. Secondary structure evaluation has been performed using JASCO software and Origin (version 2022, OriginLab Corporation, Northampton, Massachusetts, USA)^20^.
5. Raman Spectroscopy: The Raman spectra were recorded using an Alpha300 (WITec, Germany), which has a liquid nitrogen-cooled charge coupled device (CCD) detector and a spectrograph with 600 g/mm grating with resolution of 1 cm^−1^. Back-scattering geometry for spectra collection was applied with a notch filter to reject the elastic contribution. The laser emitting at 532 nm with a power of 44 mW was employed as the excitation source. (As described in the extended experimental procedures)

a. Raman spectra recording: The procedure used here was modification of a Signorelli et al., (2016)^21^ which is described in the extended experimental procedures.
b. Curve-fitting procedure: Curve fitting of the Amide I band and assignment to the structural components were performed essentially as described by Maiti et al., (2003)^22^. Curve fitting was performed via Origin (version 2022)^20^.
c. Principle component analysis (PCA): PCA was applied on Raman spectroscopy data with 10 spectra, being satisfactorily described by a principle component number 21. The correlation matrix was analyzed with Origin (version 2022)^23^ and the number of components to be considered was defined as the number required to explain at least 70% of the total variance.
6. Atomic Force Microscopy (AFM): For AFM studies, purified ΔN190TTF1 protein was diluted to 11ng/ul in 25mM Tris-Cl pH 7.5 buffer (filtered) and 10ul of this diluted protein sample was deposited on a fresh silicon wafer and dried by desiccation. AFM measurements were performed by the P9 SPM digital control platform (Solver next, NT-MDT, Russia). Images were acquired using a silicon tip (NHG-0.1 model) in tapping mode or semi-contact mode. Image processing and analysis were performed by NT-MDT Nova which was provided by the manufacturer of the machine.

## Results and Discussion

1. The results of a.a. sequence-based analysis with ProtParam showed that Full length and 190 a.a. deleted mouse TTF1 are unstable hydrophilic proteins, (instability indexes of 57.28 & 46.44) and has grand average of hydrophobicity and hydrophilicity (GRAVY) of −1.009 and −0.811 respectively. After deleting 190 a.a. theoretically we can see the instability index decreases from 57.28 to 46.66 which indicates the truncated protein is much more stable than the full length (Table 1)^24^. Proteins with hydrophobicity scores below 0 are more likely to be globular (hydrophilic), while scores above 0 are more likely to be membranous (hydrophobic) in nature^25^; therefore TTF1 is supposed to be globular and hydrophilic. Various physicochemical properties of the protein predicted by ProtParam are presented in Table 1.
2. Purification of ΔN190TTF1: ΔN190TTF1 was cloned in pMAL vector with N-terminal His tag. It was expressed in BL21-λDE3 bacterial strain and induced with 0.9 mM IPTG (Figure 2a). The induced TTF1 was extracted from lysed bacterial cell and bound with Ni-NTA column. After 10 column valume (CV) wash (as mentioned in material and methods), the protein was eluted in elution buffer (wash buffer plus 400 mM immidazole, Figure 2b). The protein at this stage was 1mg/ml and more than 95% homogenously pure (lane 5-9, Figure 2b).
3. DLS study reveals ΔN190TTF1 is mono dispersed: In order to make sure the purified protein from Ni-NTA column is soluble and homogenous, we next performed DLS to know its dispersity. DLS is used for measuring particle size ranging from nanometer to micrometer. In this procedure, by processing the time series we could find the diffusion coefficient of the molecule^26^. For this process the concentrated protein was diluted to Buffer A (15ug/ul in 25mM Tris-Cl pH 7.5). The analysis of the results obtained from size distribution by intensity (Figure 3a) and also from size distribution by number (Figure 3b) both showed that solution contained single particle size with a mean hydrodynamic radius (R_H_) of 108 nm confirming the mono-dispersive nature of the solution with pI index 0.299 (Figure 3a and b and Table 2). This evaluation allowed us to move ahead and do structural characterization of the purified protein.
4. CD spectroscopy confirms ΔN190TTF1 is a soluble and helical protein: After confirming that purified protein is monodispersed and homogenous, we wanted to know the secondary structure of the protein by performing CD spectroscopy (as described in materials and methods section). CD spectroscopy being a faster technique requiring lesser sample, plays a significant role in structural characterization and is often used as confirmatory method for secondary structure determination^27^. The far-UV CD spectrum of the ΔN190TTF1 protein (Figure 4), collected in 25 mM Tris-HCl buffer pH 7.5 at 20°C, is characterized by a sharp maxima at 192 nm and two slightly pronounced minima at 209 and 221nm. The secondary structure estimation after spectra deconvolution (as mentioned in material and methods) gave estimated percentage of α-helices (53.1%), β-sheets (6.4%) and random coils (40.4%) (Table 3), confirming predominantly the helical nature of the protein^28^. The above finding is in very much agreement with our computational ab-initio study and simulation data of human TTF1 protein^16^.
5. Raman spectroscopy of ΔN190TTF1: Raman spectroscopy is an important tool to study secondary structure of proteins and peptides. Raman spectral profile of a protein in solution is a collection of signals arising from each conformation, thus providing an “instantaneous snapshot” of the population^22,29^. Majority of the Raman spectral signals are produced by out-of-phase vibration due to C–N stretching, the C–C–N deformation and the N–H in-plane bending. The spectrum presents a complex set of peaks, and overlapping bands among which the main spectra due to aromatic amino acids and polypeptide chain can be recognized. Hence, we performed Raman spectroscopy to characterize the secondary structure and also to understand population heterogeneity of the purified TTF1. The Raman spectra of ΔN190TTF1 is shown in red colour and buffer (without protein, control) in black (Figure 5a). In the spectra, the peaks associated to Phe amino acids are located at 1000 cm^−1^. The peak around 1341 cm^−1^ are related to the Trp vibrations. At about 1250 cm^−1^, the Amide III band could be observed which occurs due to vibrations produced by in-phase combination of the N–H bending and the C–N stretching and attributes to a mixture of secondary structures in protein solution. The peak around 1550 cm^−1^ identifies Amide II band. Finally, the Amide I band, which is found within the 1620 – 1725 cm^−1^ spectral region could be observed which is caused by vibration produced mainly due to the C¼ O stretching with minor contributions. Spectra arising from Amide I modes comes from the vibrational coupling between motions of the peptide carbonyl groups organized in an ordered secondary structure through hydrogen bonds. An analysis of this band can provide detailed information on the structural conformation of the proteins in physiological conditions (Figure 5b). Since Amide I spectrum could significantly characterize secondary structure of a protein, we have described and analysed the same in detail for ΔN190TTF1 (Table 4 and Figure 5)^21,22,29,30^.
6. Curve fitting for Amide I region and deconvolution: Amide I band has been used to validate atomic structures of several proteins and also to study intrinsically disordered proteins and peptides. For these reasons we too performed curve fitting procedure for the Amide I spectrum of our protein^21,22^. In particular, the band has been deconvoluted in terms of three major components separately which are associated with the α-helix, β-sheet and random coil structures. In particular, the Amide I band has been fitted by considering a curve centred at 1650–1656 cm^−1^, which could be assigned to an α-helix, a second one at the 1664–1670 cm^−1^ region due to β-sheet, a third one at about 1680 cm^−1^, corresponding to random coils and fourth at 1640-1650 cm^−1^, attributing to the turns which connects helices. Table 4 lists the summary of corresponding fitting parameters. The contribution from the appropriate secondary structures has been connected with the integrated curve intensities fitting the Amide I band under the presumption that the Raman cross-section is the same for each conformation. The fitting curves of the Amide I Raman bands of ΔN190TTF1 protein proclaim the major area (42.44%) of the curve for α-helices (white area under red curve) that clearly shows that TTF1 mostly contains helices. The rest curve area is divided into random coils (blue curve, 21.1%) and turns (green curve, 13.03%) (Figure 5C). Amide I band full width half-maximum (FWHM) and area percentage of each curve of secondary structures for ΔN190TTF1 protein are enlisted in table 5. Deconvolution of spectra collected from CD and Raman spectroscopy shows that TTF1 is a helical protein and this finding justifies its DNA binding activity. The above finding is in agreement with our computational model for full length TTF1 as well^16^. Previous literatures reporting protein DNA atomic structures have shown that the DNA is stabilized between the helices of those proteins^2,31^.
7. Principle Component analysis of the Amide 1 region: Additionally, Amide I region (1600-1700 cm^−1^) have been analysed through the PCA technique which is a multivariate statistical analysis method that allows one to reduce the multi-dimensionality of the dataset by recollecting the characteristics of the original ensemble, primarily contributing to its quantitative variance. Concisely, a limited number of variables, known as principal components (PCs), contain most of the spectral information and they can be divided into patterns, scores, and loadings. Figure 6a shows the PCA scores of PC1 vs PC2 component for Amide I band of ΔN190TTF1 (experimental, black dots), and buffer (control, red dots). PC1 and PC2 components of Amide I band of ΔN190TTF1 combined shows 81% of the total variance. The plot shown in Figure 6a represents the distribution of the spectra for a given PC and reveals the relationship existing among the samples. While loading plot shown in Figure 6b, represents how much an original variable, or range of variables, influences a given PC for a particular defined region. Hence, PCA is particularly effective for categorizing Raman spectra that are otherwise hardly distinguishable^21,32^. Along the PC1 axis, two separate clusters can be seen. Most of the scores of ΔN190TTF1 are located in the positive portion of plot along PC1 with a spread along both the axes, while the control scores also display both negative and positive PC1 values with a large variability along PC2. The PC2 loading plot, providing complementary information to PC1, shows a significant variation in 1600-1700 cm^−1^ region. Variation specifically corresponding to tyrosine peak (1613-1616 cm^−1^) could be observed, that occurs mainly due to change in strength of hydrogen bond between water and tyrosine molecule. Downshifting is observed in the band frequency in region 1670-1685 cm^−1^ when helices form coils. A high spectral variance also suggests the presence of α-helices (1646 cm^−1^), turns (1635 cm^−1^) and random coils (1681 cm^−1^) ^30^, as previously observed (Figure 5b and Figure 6b). PCA strengthens our Raman spectral data via introducing theoretical statistical calculation to show the quantitative variability in buffer components and the existence of ΔN190TTF1protein in optimized buffer.
8. Atomic Force Microscopy: AFM imaging of single protein molecules are used to quantitatively evaluate their dimensions and is quite useful to determine its geometrics per unit area of interface, which eventually fascilitates the determination of size of the molecule^26^. For this, a area of 10 um X 10 um was scanned and topography of the surface was determine as shown in figure 7. After the calculations (as shown in micrograph of Figure 7a), average size of the molecule was determined to be 0.094 um (94nm, Figure 7c), which is almost consistent with the DLS sizing experiment results as reported in previous section (Figure 3a). DLS data represented the hydrodynamic size (R_H_) while AFM imaging shows the physical diameter of the particle (therefore the differences in size are natural)^33^. In figure 7b the differences in baseline of top and bottom Z-axis indicate the dimension of protein molecules. Hence, AFM study confirmed the homogeneity of the purification and revealed molecular size of the protein as well.

**Figure 2:**
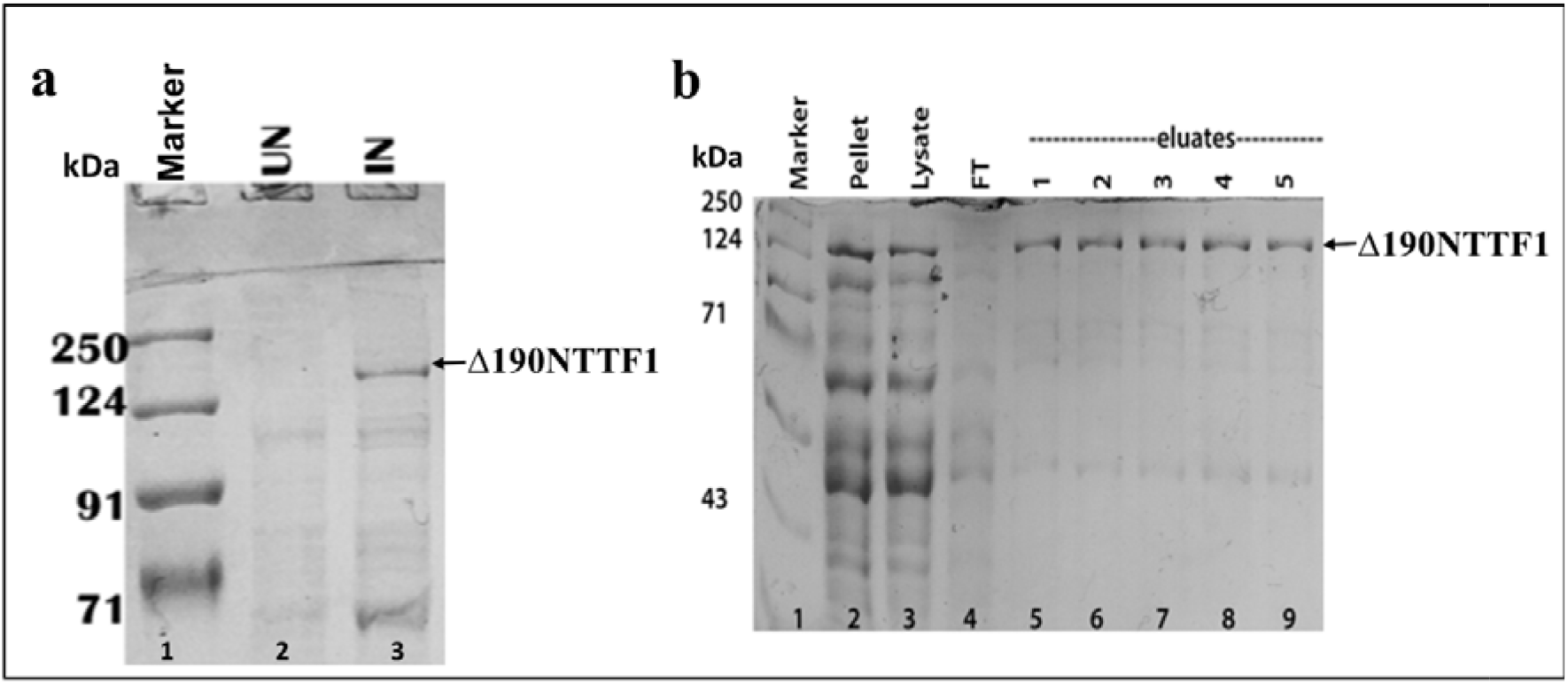
**a)** Induction profile of ΔN190TTF1 protein; “In” represents induced (lanes 3) and “Un” represents un-induced expression profile of the same (lane 2); **b)** Purification profile of ΔN190TTF1 protein; Lane 2 and 3 shows protein profile of the pellet and lysate respectively and lane 5-9 shows eluted TTF1.

**Figure 3.**
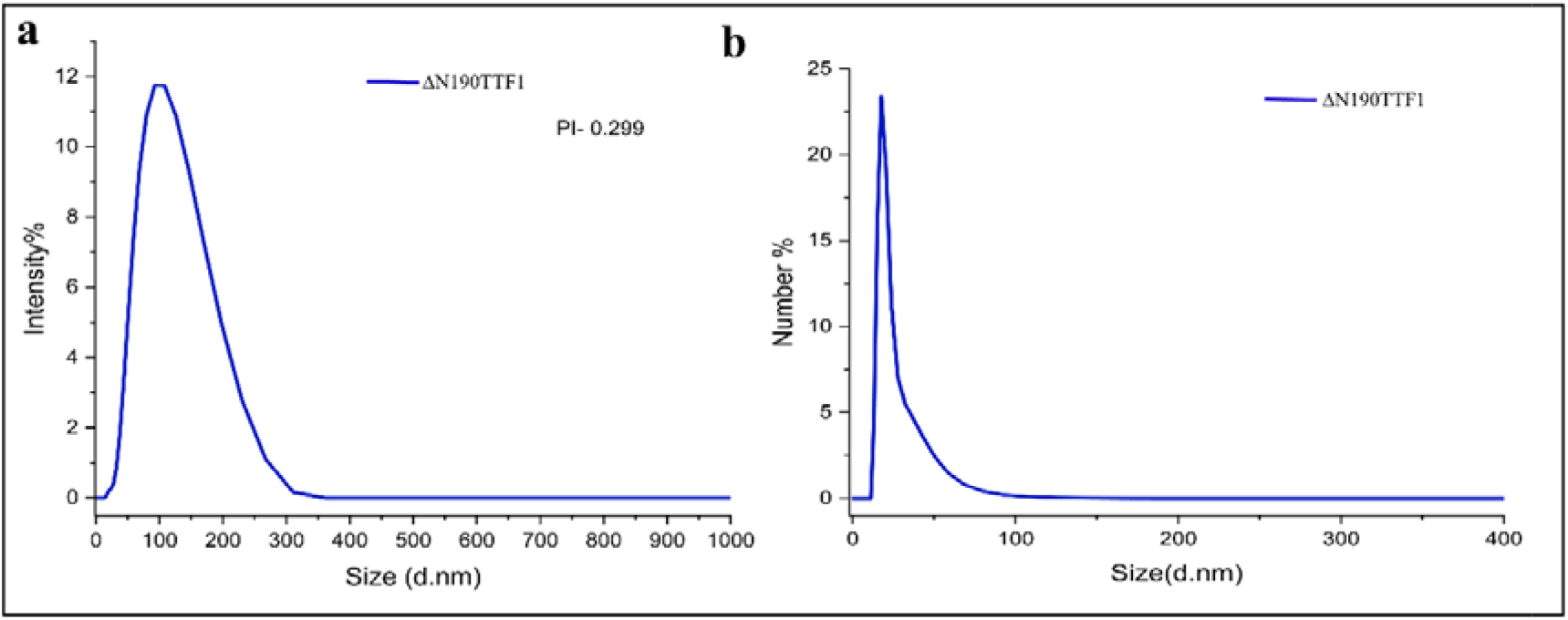
DLS analysis plots of purified ΔN190TTF1 protein, **a)** size distribution by intensity plot; **b)** size distribution by number plot.

**Figure 4.**
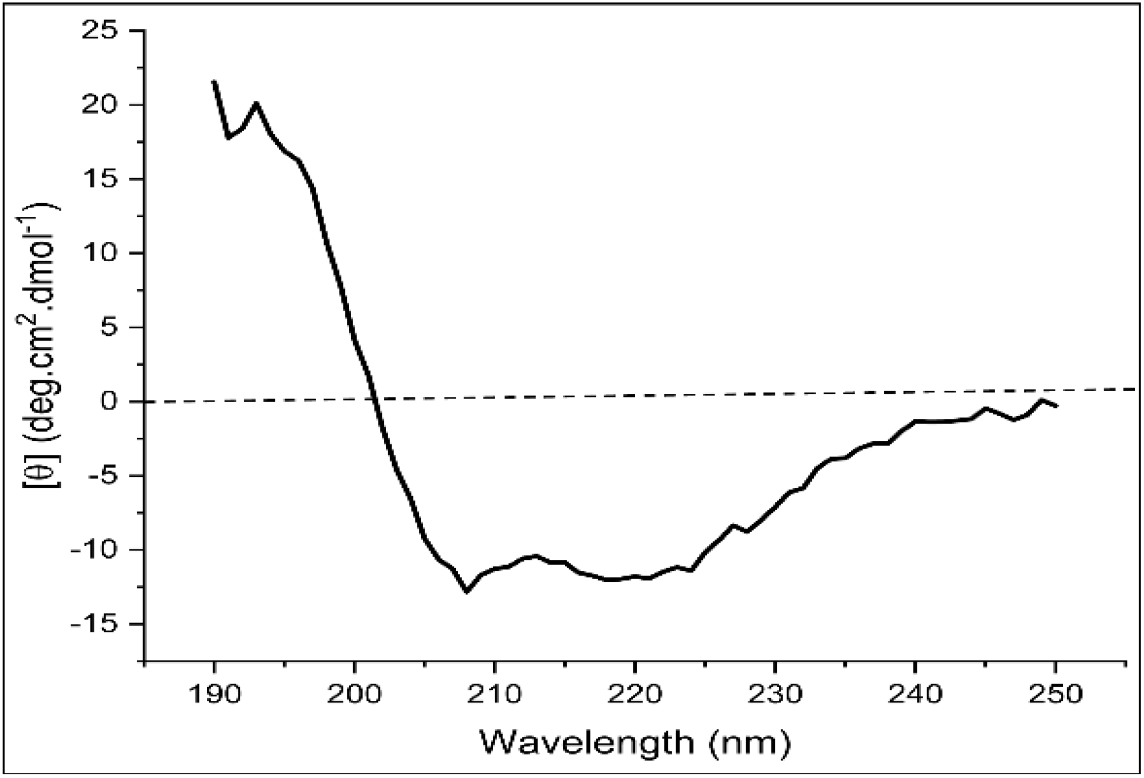
CD spectra of ÄN190TTF1 protein.

**Figure 5:**
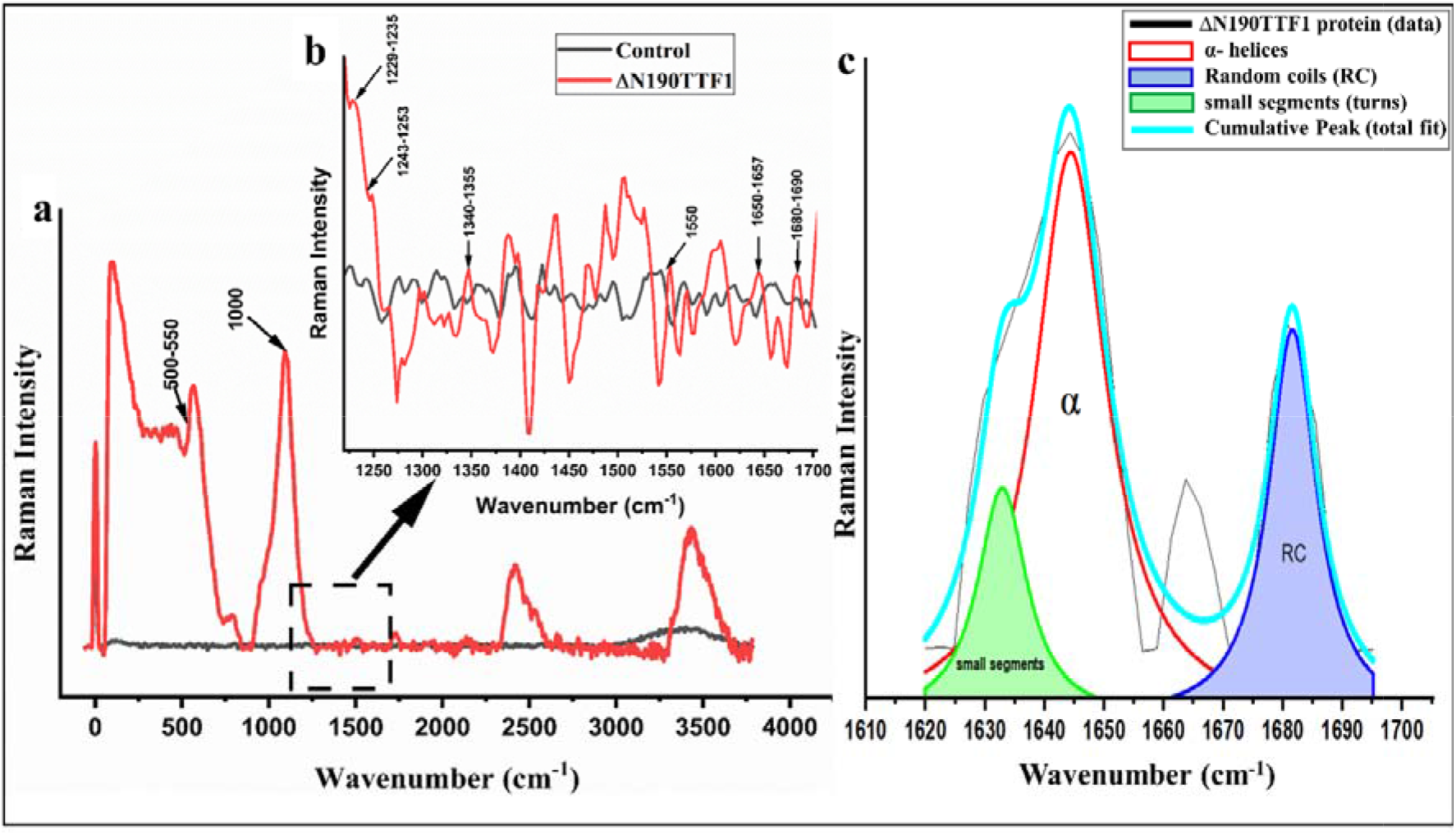
**a)** Raman spectra of ΔN190TTF1 protein; **b)** Raman spectra for 1200-1700 cm^−1^ region (magnified). TTF1 spectrum is shown in red while that of control (buffer without protein) is shown in black; **c)** Raman spectra after curve fitting for the experimental Amide I data.

**Figure 6.**
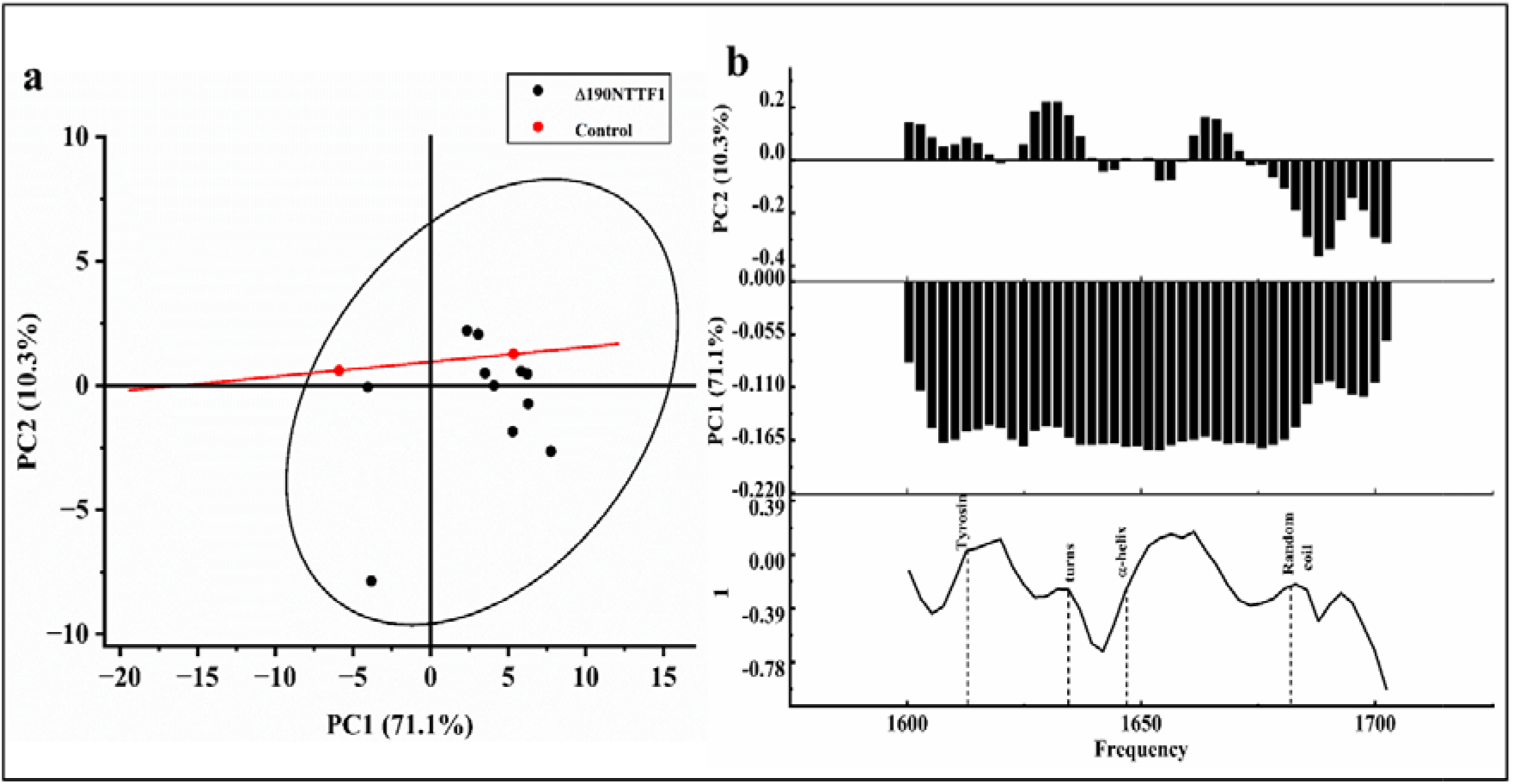
**a)** Two-dimensional score plot PC1 verses PC2 of the Raman spectra for ΔN190TTF1 (black dots) and control (buffer; red dots) performed on Amide I band. Protein group are illustrated as ellipse and control group as line; **b)** one dimensional loading plot versus frequency (PC2 10.3% and PC1 71.1% of total variance).

**Figure 7.**
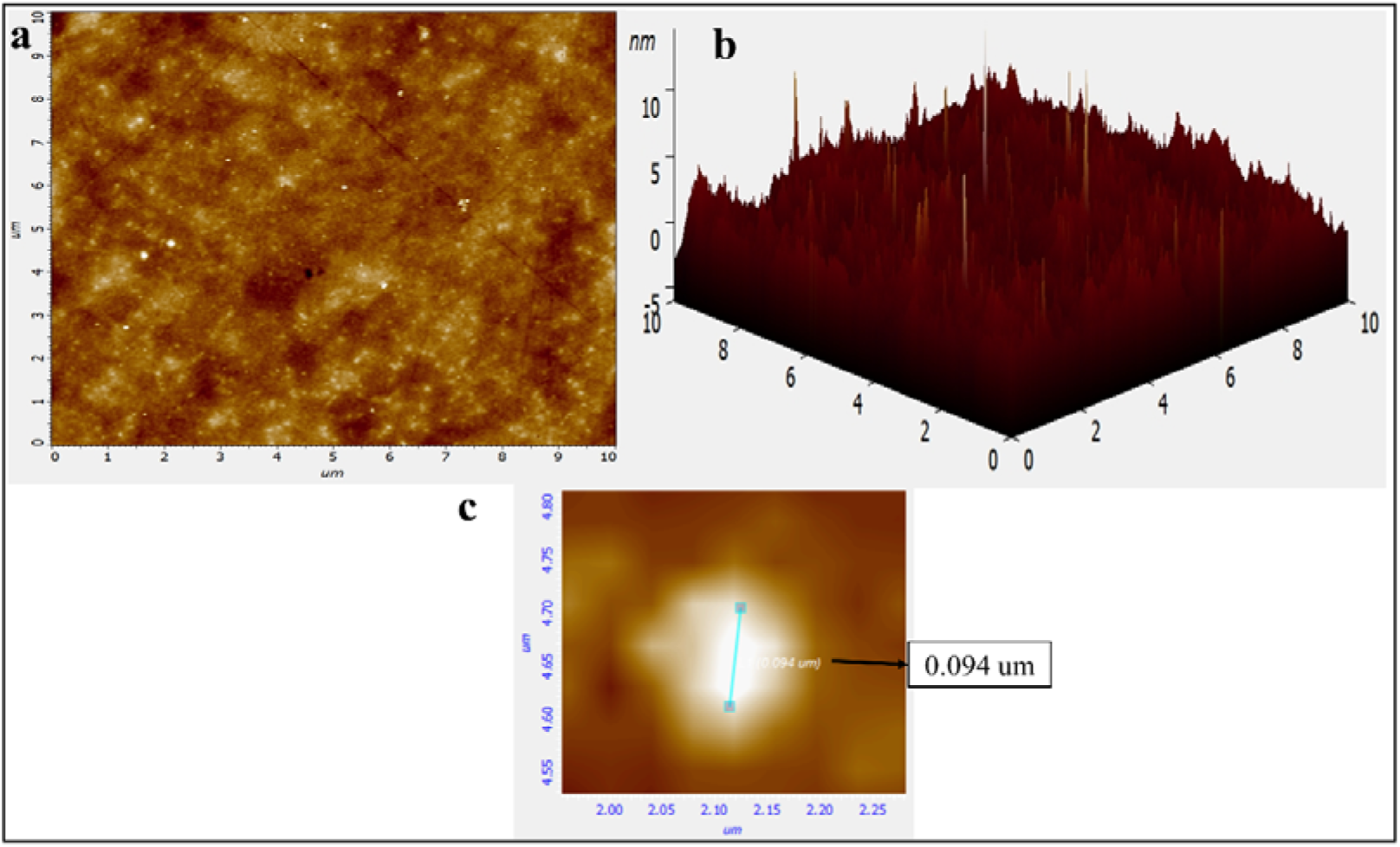
AFM micrograph 10 × 10 um of ΔN190TTF1 protein, **a)** ΔN190TTF1 2D image; **b)** ΔN190TTF1 3D image; **c)** Shows a ΔN190TTF1 protein molecule under higher magnification.

**Table 1:** Physicochemical properties of mouseTTF1, as determined with ProtParam.

| Physicochemical Properties | Values |  |
| --- | --- | --- |
|  | Full length mouse TTF1 | 190 a.a. deleted mouse TTF1 |
| Number of residues | 859 | 669 |
| Molecular formula | C<sub>4262</sub>H<sub>6916</sub>N<sub>1256</sub>O<sub>1333</sub>S<sub>20</sub> | C<sub>3317</sub>H<sub>5323</sub>N<sub>951</sub>O<sub>1043</sub>S<sub>17</sub> |
| Molecular weight | 97722.61 Da | 75758.52 Da |
| Theoretical pI | 9.41 | 7.93 |
| Instability index | 57.28 | 46.66 |
| Aliphatic index | 65.42 | 73.06 |
| Total number of negatively charged residues (D+E) | 133 | 112 |
| Total number of positively charged residues (R+K) | 166 | 114 |
| Grand average of hydrophobicity (GRAVY) | -1.009 | -0.811 |
| Estimated half-life (mammalian reticulocyte, *in vitro*) | 30 hours | 30 hours |

**Table 2.**
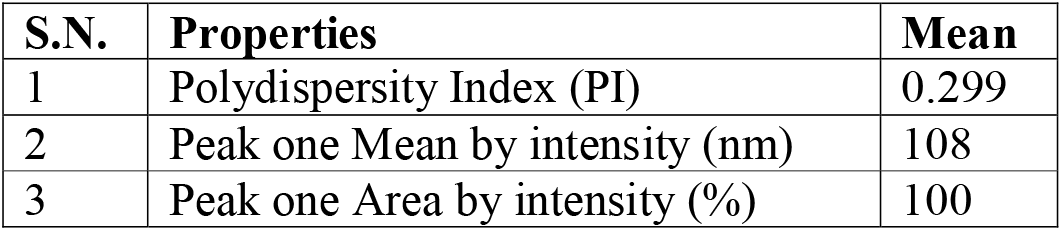
Properties of ΔN190TTF1 protein in 25mM Tris pH 7.5 buffer after DLS analysis.

| S.N. | Properties | Mean |
| --- | --- | --- |
| 1 | Polydispersity Index (PI) | 0.299 |
| 2 | Peak one Mean by intensity (nm) | 108 |
| 3 | Peak one Area by intensity (%) | 100 |

**Table 3.** Secondary structure percentage estimated after deconvolution of CD spectra.

| S.N. | Sec. Structure | Percentage (%) |
| --- | --- | --- |
| 1 | α-helix | 53.1 |
| 2 | β-sheet | 6.4 |
| 3 | Random coil | 40.4 |

**Table 4:** Raman bands of ΔN190TTF1 protein spectra with the assignment^29,30^.

| Source | Frequency (Raman band position $\text{cm}^{-1}$) | Assignment |
| --- | --- | --- |
| Amide I | 1650-1657 | $\alpha$ helices |
|  | 1680-1690 | Turn |
| Amide II | 1550 | Parallel/anti parallel $\beta$ sheet |
| Amide III | 1243-1253 | Random coil |
| | 1229-1235 | $\beta$ sheet |
| Tryptophan | 1340-1355 | Relative ratio is indicative of hydrophobicity |
| Phenylalanine | 1000 | CN stretching |
| Disulfide | 500-550 | S-S bond stretching |

**Table 5:** Results from the curve fitting procedure of the Amide I band of ΔN190TTF1. (FWHM: Full width at half maximum)

| Secondary structure | Frequency (cm<sup>-1</sup>) | Area (%) | FWHM (cm<sup>-1</sup>) |
| --- | --- | --- | --- |
| α-helix | 1646 | 42.44 | 15.95 |
| Small segments (turns) | 1635 | 13.03 | 11.75 |
| Random coil | 1681 | 21.34 | 11.44 |

## Conclusion

Even though TTF1 being an essential multifunctional protein, almost nothing is known about its biophysical character or structure. The only study revealing its structural model (computationally) is reported from our lab^16^. Hence, it’s important to understand the biophysical characters of this protein in order to engineer this protein for structural and mechanistic studies. Towards achieving the above objectives, we first assayed the homogeneity of purified protein and then characterized the secondary structure. The purified protein profile on gel, DLS and AFM data confirmed the homogeneity and stability (in optimized buffer) of the preparation, which gave us confidence to proceed further for structural characterization. CD and Raman spectroscopy established that ΔN190TTF1 is a helical protein. The deconvolution of the Amide I band, remarkably sensitive to the α-helix, ß-sheets and random coil structures, has allowed us to assess the main structural motifs of these proteins. In particular, a careful fitting procedure of this band has revealed that about ~50% of native ΔN190TTF1 displays α-helices, while the rest is almost equally shared between random coil and small segment structures. The AFM imaging furnished reliable information about size of single ΔN190TTF1 molecule to be ~ 94 nm. Our lab is aggressively working to solve the physical structure of this protein by cryo-electron microscopy and our current finding in this article infuses confidence in us that we are moving in right direction. Knowing the biophysical and structural properties of this essential protein, will significantly help researchers to know the effect of mutation and its role in diseases like Cockayne syndrome B and various others to be discovered yet.

## Acknowledgement

The authors are thankful to the Coordinator, School of Biotechnology, and Director Institute of Science, Banaras Hindu University for providing space and facilities to conduct the research. We also thankful to Central Discovery Centre (CDC) and Sophisticated Analytical and Technical Help Institutes (SATHI), BHU for the accessibility of Spectropolarimeter and Raman Spectroscopy instruments. We also thank department of Chemistry, Institute of science, BHU for DLS and AFM instrument facility and expertise advice regarding the same. Further, we are thankful to Department of Biotechnology (DBT), Govt. of India for funding SKS with RLS grant (BT/RLF/Re-entry/43/2016) and JRF fellowship to KT. We are also thankful to IoE, BHU and University Grant Commission for providing funds to SKS. We also thank CSIR for funding the JRF fellowship to GS.

## Supplemental Information

### Extended experimental Procedure

1. **Expression and purification of recombinant TTF1**: Codon optimized ΔN190TTF1 cloned and subcloned in pMAL vector. Then the recombinant TTF1 was amplified in *Escherichia coli (E.coli*) strain DH5α and transformed in *E.coli* strain BL21-λDE3. Exponentially growing transformed BL21-λDE3 was induced with 0.9mM IPTG at OD_600_= 0.8 and the temperature was shifted to 18°C overnight. Bacterial culture was harvested and lysed in Lysis buffer [25mM Tris (pH-7.5), 500mM KCl, 10% (w/v) glycerol, 9mM ß-mercaptoethanol, 5mg/ml Lysozyme and protease inhibitor cocktail]. The cells were ultra-sonicated on ice using 15 s pulse at 30 amplitude for 3 mins with 1 min interval between each pulse. The clear lysate was mixed with prepared Ni-NTA beads and kept on rotary shaker for 1 hr at 4°C for binding with the desired protein, which was then loaded onto the silica column and allowed to settle by gravity. Washed column thrice with 10 column volume wash buffer (25mM Tris (pH-7.5), 500mM KCl, 10% (w/v) glycerol, 9mM β-mercaptoethanol). The fusion protein was then eluted with an elution buffer [wash buffer with 400mM imidazole]. 8% SDS-PAGE was run at each step of the whole purification process to observe induction and purification.
2. **Raman spectra recording**: The Raman spectra of the ΔN190TTF1 protein measurements were collected using a 50 objective with a numerical aperture (NA) ¼ 0.6 (laser spot diameter reaching the sample was about 1 mm). Raman experiments were performed on sample (2ug/ml) drops deposited on optical glass slide with cavity, at room temperature. 5 Raman spectra for each sample were collected from different points of the drops, each one with an acquisition time of 10 s. Furthermore, no variation in the band spectral features (i.e., changes in shape or in vibrational frequency) was observed during the acquisition time. All Raman spectra were baseline corrected after being vector normalized across the whole wavenumber range. A linear baseline correction was used in the majority of cases. After comparing all recorded spectra and confirming the spectral profile, spectra with high signal-to-noise ratio were chosen. The spectra was plotted and analyzed by Origin, version 2022 (**OriginLab** Corporation, Northampton, Massachusetts, USA).

